# CRISPR adaptation in *Streptococcus thermophilus* can be driven by phage environmental DNA

**DOI:** 10.1101/2024.06.13.598888

**Authors:** FR Croteau, J Tran, AP Hynes

## Abstract

The CRISPR-Cas system is a bacterial adaptative immune system which protects against infection by phages: viruses that infect bacteria. To develop immunity, bacteria integrate spacers — fragments of the invading nucleic acids — into their CRISPR array to serve as the basis for sequence-targeted DNA cleavage. However, upon infection, phages quickly take over the metabolism of the bacteria, leaving little time for the bacteria to acquire new spacers, transcribe them and use them to cut the invading DNA. To develop CRISPR immunity, bacteria must be safely exposed to phage DNA. Phage infection releases eDNA which could be involved in the development of CRISPR immunity. Using *S. thermophilus* and phages 2972 and 858 as a model for CRISPR immunity, we show that eDNA is crucial to the development of optimal CRISPR immunity, as generation of phage-immune bacterial colonies decrease with eDNA digestion. Furthermore, it is phage eDNA specifically that impacts CRISPR immunity since its addition increases the generation of phage-immune colonies. We also show that the effect of eDNA is phage-specific, sequence specific and can even be traced to a region of the genome covering the early-expressed genes which differ between phages 2972 and 858. However, we also show that eDNA is not used as a source of genetic information for spacer acquisition. This suggests that the effect of eDNA involves a new mechanism of phage resistance. Moreover, the effect of eDNA is highly dependent on environmental conditions as variation in media suppliers are sufficient to interfere with this effect. These results link environmental conditions, specifically eDNA, to the CRISPR-Cas system, providing a better understanding of the context of the emergence of CRISPR immunity and could inform our understanding of the mechanisms through which bacteria detect the presence of phages before infection.

## Introduction

The CRISPR-Cas system, composed of an array of clustered regularly interspaced short palindromic repeats (CRISPR) and associated *Cas* genes, is a bacterial adaptive immune system (1) that protects against foreign DNA, including bacteriophages. The array is interspaced with sequences called spacers, matching invading genetic elements such as phages or plasmids (2). These spacers form the basis of the immunity as they are transcribed and processed into crRNAs (3) which then guide endonucleases to induce site-specific cleavage of any matching invading sequences (4). The development of new CRISPR-based immunity, through the acquisition of new spacers, has been observed both in laboratory and natural settings (1). Phage DNA is indubitably a source of spacer sequences, and while advances have been made in the identification of many of the molecular steps of spacer acquisition (5–12), none satisfactorily explain the ‘paradox of timing’ inherent in this immunity.

The paradox is conceptually simple: the CRISPR-Cas system requires exposure to phage DNA to provide immunity. In the absence of resistance, intracellular exposure to phage DNA will almost invariably lead to phage replication and the death of the cell. Even transfection of purified or synthetic phage genomes into a bacterial cell leads to production of phage particles (13). To combat this, the CRISPR-Cas system rapidly cleaves matching invading DNA, with some cleaved DNA being detected as quickly as 2 min after infection (4), by using pre-existing complexes of guide RNAs and Cas endonucleases (3, 14–16). The short time frame of this response is further demonstrated by the speed of the phage anti-CRISPR response. While phage tools to block CRISPRs range from dedicated anti-CRISPR proteins (17) to nucleus-like protective compartments (18), they are active within minutes of infection (19), or, in the case of nucleus-like structures, concurrently with DNA entry. Given the short time scale on which the battle for the fate of the cell is fought: how are CRISPR-naïve cells meant to acquire a new spacer, transcribe it into RNA, mature the crRNAs and then find and cleave the invading sequences before the phage has hijacked the system, or irreparably damaged the cell?

Despite this timing issue, the CRISPR-Cas system manages to acquire new spacers from foreign sources of DNA – even in the absence of other involved mechanisms of resistance, models of CRISPR adaptation such as *Pseudomonas* and *Streptococcus* regularly yield CRISPR-immune survivors of phage challenges at rates of ∼1/10^6^ (20). These rates can be improved by synergy with other defence mechanisms (21–24), or by pre-existing partial spacer matches able to ‘prime’ the immunity (25–27) – but neither of these directly address the problem of a naïve cell, dependent on the CRISPR system, in which such an adaptive immune system must have been selected for. In some cases, machinery essential to the CRISPR adaptation process itself - RecBDC/AddAB complexes - degrade invading phage DNA and generate DNA free ends suitable for spacer acquisition (8). However, if this were an effective means of inactivating the phage to generate new spacers, the CRISPR system would be somewhat redundant – depending entirely on the success of an innate immune system.

One solution to this conundrum was demonstrated in 2014 when defective phages, capable of DNA injection but otherwise not capable of completing the rest of the infection, were shown to drive CRISPR adaptation in *S. thermophilus* (28). This demonstrated one way in which bacteria could be safely exposed to foreign DNA that was no longer capable of leading to cell death, thus removing the time pressure seen in successful infection. This is somewhat analogous to vaccination in enabling an adaptive immune system to overcome a highly virulent virus such as rabies, where the immune system would normally have no opportunity for a repeat exposure to the virus. However, the proposed model assumed that lysate contained 10% inherently defective phages which are responsible for >96% of all acquisition events. The authors acknowledged that this is untestable using current techniques. This leaves the door open for other possible ways for the bacteria to be safely exposed to phage DNA.

Bacteria get exposed to foreign DNA in one of three ways, transformation, conjugation and transduction. The acquisition of spacers from DNA injected by defective phages is similar to transduction and spacers are readily acquired from plasmids (29)and making it likely that conjugation could lead to acquisition. However, transformation has yet to be linked to the development of phage immunity despite the readily available phage genetic material in the environment. In the context of phage infection, phage-induced lysis will cause the release of any number of remaining intracellular components including bacterial and phage proteins, small molecules such as nucleotides, bacterial DNA and unencapsidated phage DNA (30). This phage environmental DNA (eDNA) contains the genetic information necessary for CRISPR adaptation and defence and can be taken up by naturally competent bacteria through transformation. Moreover, before being taken up, the eDNA is cut into smaller pieces (31) and one of the strands of the eDNA is degraded so that only a single strand of DNA enters the cell (32, 33). In the context of exposure to phage genetic material, this would make transformed DNA safe for the bacteria. We hypothesized that eDNA released as part of phage lysis could be involved in the development of CRISPR immunity.

Using the model of CRISPR-Cas adaptation, *Streptococcus thermophilus* DGCC7710, we show that eDNA is an important contributor to the generation of new CRISPR immunity, that this effect is not only specific to the phage – requiring the environmental DNA to match the challenging phage, but surprisingly does not occur by providing a source of genetic material for spacer acquisition.

## Results

### Phage lysates contain both bacterial and phage eDNA

To confirm the presence of eDNA in phage lysates, we measured the concentration of both bacterial and phage eDNA using qPCR with species-specific primers in lysates of phage 2972 and phage 858. Bacterial eDNA was detected in phage 2972 lysates at 0.13(3) ng·µl^-1^ (n=9) and in phage 858 lysates at 0.157(3) ng^·µl-1^ (n=4) for an overall average of 0.14(2) ng·µl^-1^ (n=13).

To detect phage eDNA, we had to assume the high temperature ((95°C) of the PCR could result in the release of packaged phage DNA. In order to calculate the concentration of phage eDNA present before qPCR, we subtracted the value of phage eDNA measured in a lysate treated with DNase, which contains eDNA released during the qPCR and during the inactivation of the DNase, from the value measured from the same lysate treated with DNase buffer only, which contains the original eDNA as well as eDNA released during both the qPCR and the inactivation of DNase. Phage eDNA was detected in phage 2972 lysates at 0.5(1) ng·µl^-1^ (n=9) and in phage 858 lysates at 0.6(2) ng·µl-^1^ (n=6) for an overall average of 0.5(1) ng^·µl-1^ (n=15). This corresponds to 10.40e7 and 1.65e7 genomes per microliter for phages 2972 and 858 respectively. In our lysates free phage genomes outnumbered infectious particles 100:1, enough for more than 50 free-floating phage genomes per bacterial CFU in a typical BIM assay. There was no significant difference in the eDNA value measured across the lysates from the two different phage species (Welch’s Anova, p>0.05). We also plotted the calculated eDNA concentration in each lysate as a function of phage titre and determined through linear regression analysis that there was no significant correlation between phage titre and phage eDNA concentration for either phage (Supplementary Figure 1).

To establish the efficacy of the DNase treatment in phage lysate, we measured bacterial eDNA concentration in DNase treated lysates. After digestion, the average concentration of bacterial eDNA in lysates dropped to 0.00(2) ng^·µl-1^ (n=13), which is a significant reduction in measurable eDNA level when compared to the untreated condition (p<0.001, Welch’s ANOVA, Games-Howell test).

### DNase treatment reduces BIM generation

Having confirmed of the presence of phage eDNA in lysates and the efficacy of DNase in digesting it, we tested the effect of eDNA digestion on the generation of CRISPR-mediated immunity by comparing the number of bacteriophage insensitive mutant (BIM) colonies obtained after a phage challenge with untreated or DNase-treated lysates, using lysates treated with DNase buffer only as a control. The challenges we performed at a MOI between 0.3 and 0.4 since BIM generation tends to peak in those conditions (20). In challenges with phage 2972, the DNase treated lysates yielded 40(5)% fewer BIMs than the untreated control condition (Welch’s Anova, Games-Howell test, p<0.001). The untreated condition produced a number of BIMs consistent with the expected average acquisition rate of ∼1 in 1e6 (20). The DNase buffer condition did not result in a significant difference from the control (Figure 1A). This experiment was repeated using phage 858 for the challenge and yielded similar results with DNase treated lysates resulting in 32(7)% fewer BIMs than the untreated control condition (p<0.05, Games-Howell test) (Figure 1B).

**Figure 1:**
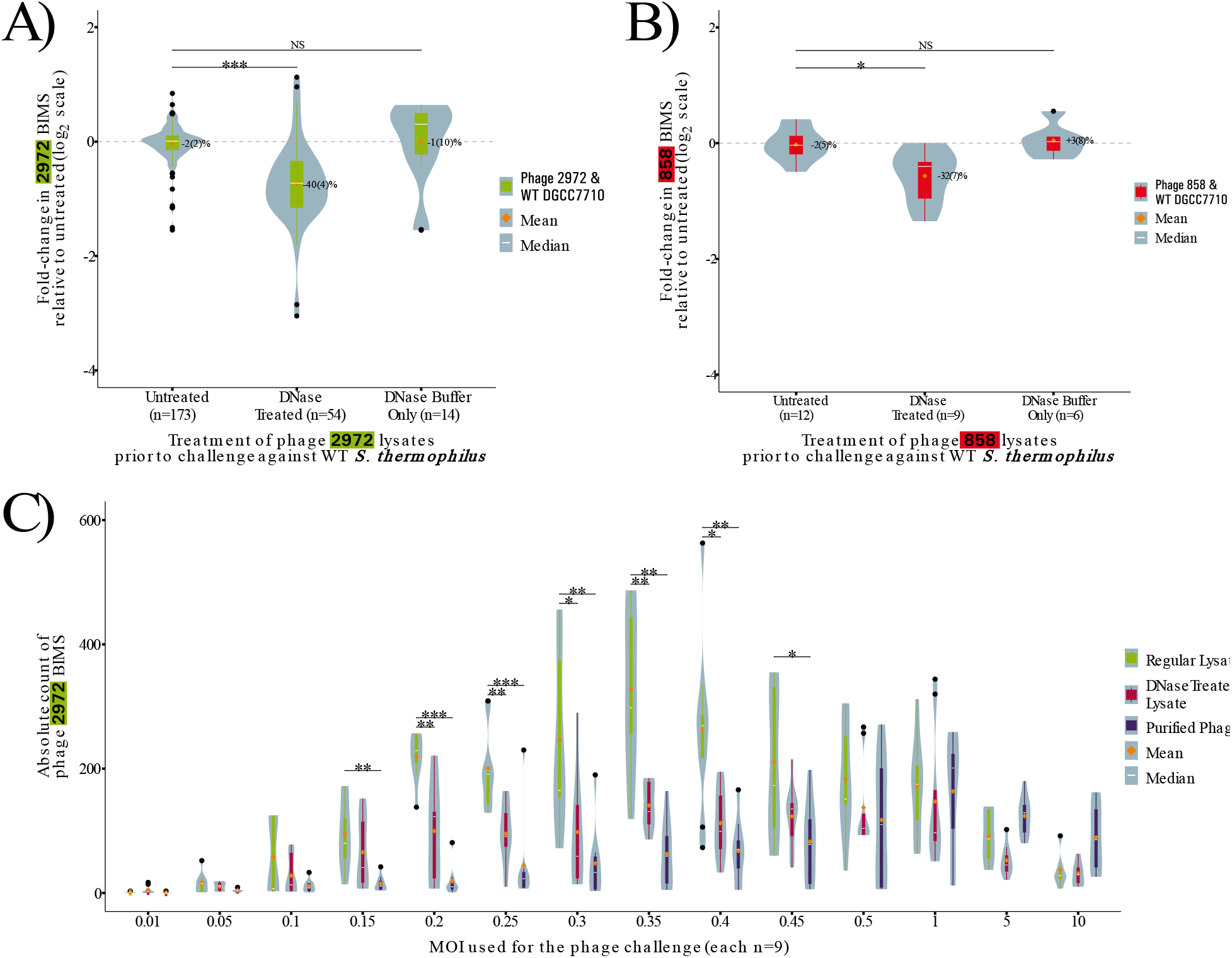
BIM generation with DNase treated lysate and purified phage. A-B) BIMs obtained in challenges against phage 2972 (A) and phage 858 (B) at MOIs between 0.15 and 0.45. BIMs are reported as fold-change relative to the baseline. C) BIMs obtained in challenges against phage 2972 at varying MOIs. BIMs are reported as absolute counts for each condition in either untreated lysates, DNase treated lysates or purified phages. In all cases, violin plots represent the probability density curve of the distribution, boxplots represent the first and third quartile of the distribution (box), the minimum and maximum (whiskers), the median (white line) as well as any outliers (black dots). The mean of each distribution is represented by an orange diamond. Significance is determined by pairwise comparison with the untreated condition using Welch’s Anova followed by Games-Howell post-hoc pairwise comparison test.

We attempted to recapitulate the phenotype by digesting eDNA with various restriction enzymes. However, digestion by restriction enzymes was not sufficient to cause a significant effect on BIM generation (Supplementary Figure 2). We also verified that the DNase treatment did not affect the concentration of plaque forming units (PFU) in phage 2972 and found that the variation in titre as calculated by full plate PFU count assays was -1(1)%, which is not significant (Tukey’s HSD test, P>0.05) (Supplementary Figure 3). Since there is no effect caused by treatment with DNase buffer only, the reduction in BIMs is therefore due to the degradation of eDNA by DNase. This shows that eDNA plays a role in the acquisition of resistance against phages.

### The effect of eDNA is MOI-dependent

To verify if this effect of DNase was universal across multiple MOIs, we performed similar challenges at multiple MOIs ranging from 0.01 to 10. We performed these challenges using untreated lysates, DNase treated lysates as well as purified phage solutions obtained by PEG precipitation and ultracentrifugation to remove not only eDNA but also all other lysate components. In untreated lysates, BIM generation peaked at a MOI of 0.35, confirming prior results indicating this MOI range as optimal for generation of CRISPR immunity (20). The DNase treatment had a significant effect on BIM generation at MOIs between 0.2 and 0.4 (p<0.05, Welch’s ANOVA, Games-Howell test) (Figure 1C). The purified phage solution yielded significantly fewer BIMs than the control at MOIs between 0.15 and 0.45 (p<0.05, Welch’s ANOVA, Games-Howell test) (Figure 1C). This shows that the effect of eDNA is MOI-dependent as at higher MOI all conditions yielded similar amounts of BIMs, implying that, at lower MOIs, uninfected cells benefit from the “first wave” of lysis, supporting our hypothesis that eDNA is involved in CRISPR adaptation.

To determine whether eDNA was solely responsible for this effect, we supplemented purified phage with an excess of phage eDNA in an attempt to restore the phenotype. However, the supplementation of phage eDNA did not increase BIM generation when compared to untreated purified phage (Supplementary Figure 4). This indicates that while eDNA is necessary for “optimal” BIM generation, it is not sufficient. The effect likely requires other cofactors to be effective. We tested if the addition of the ComS competence inducing peptide alongside eDNA could be sufficient to restore the phenotype since this short signal peptide has been shown to induce natural competence in *S. thermophilus* (34) but the addition of the peptide alongside eDNA had no effect (Supplementary Figure 4).

### Competence is likely necessary for the effect of eDNA

In order to test if eDNA was taken up by the bacteria to provide its immunogenic effect, we generated a scarless deletion for the *comEC* gene which encodes the ComEC channel protein, a necessary part of natural competence in gram positive bacteria (35). The sensitivity of the Δ*comEC* strain to both phages used was tested using an efficiency of plaquing (EOP) assay, yielding an EOP of 0.997 for phage 2972 and 1.029 for phage 858, confirming that the deletion of the *comEC* gene did not impact the sensitivity to either phage. In the Δ*comEC* strain the absolute number of BIMs generated was significantly reduced when compared to the WT strain (p<0.05, Welch’s Anova, Games-Howell test) (Supplementary Figure 5A). Moreover, the effect of DNase treatment of lysates on BIM generation was lost when the Δ*comEC* strain was used for phage challenges (Supplementary Figure 5B). We could not produce a complementation strain to confirm these results were due only to the *comEC* deletion but believe the loss of the effect of DNase in the ΔcomEC strain indicates that eDNA is taken up through competence in order to affect CRISPR adaptation.

### Supplementation of phage eDNA can increase BIM generation

In order to determine which properties of eDNA were responsible for its immunogenic phenotype, we tested if we could bolster BIM generation by supplementing purified DNA to phage lysates. Ideally, this experiment would have been performed in a DNase-treated lysate, but we were unable to inactivate or sequester the DNase without affecting the viability of infectiousness of the phages. First, we supplemented to a final concentration of 10 ng·μl^-1^ whole genomic DNA from WT *S. thermophilus*, from a BIM *S. thermophilus* strain with a spacer targeting phage 2972, from phage 2972 itself and from the related phage 858. This was to verify if the effect of eDNA was due to any DNA molecule being detected, if bacteria could use pre-adapted CRISPR arrays to expand their own through recombination or if phage DNA was required to serve as early warning. Across these conditions, only the supplementation of DNA from phage 2972 resulted in a significant increase (p<0.001, Welch’s Anova, Games-Howell test), with 33(8)% more BIMs generated (Figure 2A). This confirms that phage DNA is driving the effect of eDNA on BIM generation. Moreover, despite sharing most of its genome with phage 2972, the supplementation of phage 858 DNA did not increase BIM generation upon selection with 2972, potentially due to factors unique to either phage’s genome – such as unknown modification systems.

**Figure 2:**
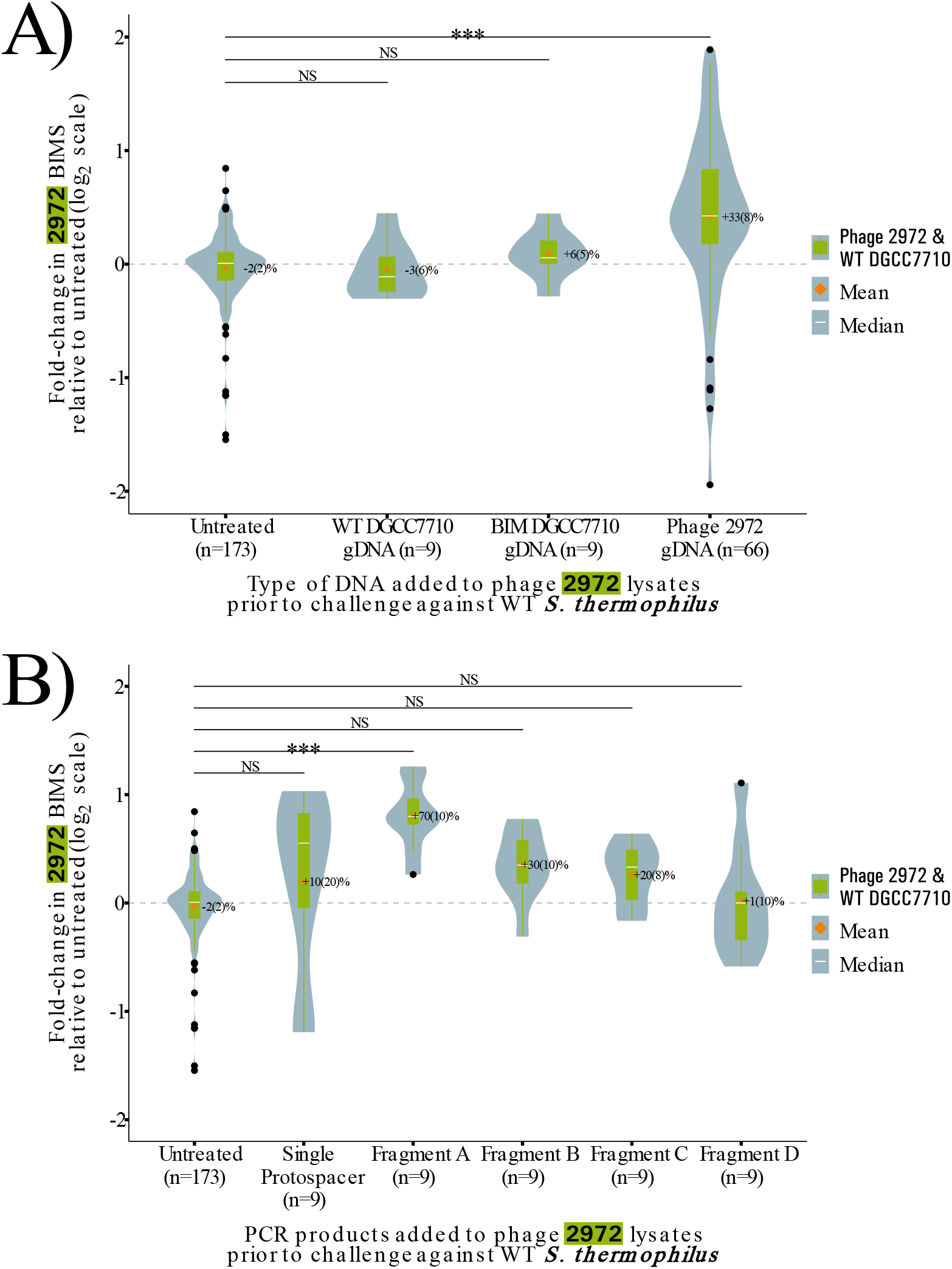
BIM generation with lysates supplemented with purified DNA. A) BIMs obtained in challenges with phage 2972 lysates supplemented with 10 ng·μl^-1^ of DNA from various sources B) BIMs obtained in challenges with phage 2972 lysates supplemented with 10 ng·μl^-1^ of PCR amplicons. In all cases, BIMs are reported as fold-change relative to the baseline, violin plots represent the probability density curve of the distribution, boxplots represent the first and third quartile of the distribution (box), the minimum and maximum (whiskers), the median (white line) as well as any outliers (black dots). The mean of each distribution is represented by an orange diamond. Significance is determined by pairwise comparison with the untreated condition using Welch’s Anova followed by Games-Howell post-hoc pairwise comparison test.

### Specific sequences of phage eDNA are responsible for the increase in BIM generation

To further examine the mechanism through which phage eDNA contributes to CRISPR adaptation and to verify if epigenetic factors are responsible for lack of effect observed when supplementing DNA from phage 858, we designed PCR amplicons of varying sizes across the genome of phage 2972 to see if the effect observed with the supplementation of phage 2972 gDNA could be recapitulated with PCR generated DNA. We identified phage genomic regions containing similar number of protospacers, the regions next to the protospacer-adjacent motifs that allow for CRISPR acquisition (2), that could be potentially targeted by the CRISPR 1 (CR1) array of *S. thermophilus*. This array was chosen since it is responsible for upwards of 90% of all acquisition events(36).

We designed five amplicons; two amplicons of approximately 600 bp containing 7 protospacers, two of approximately 1000 bp containing 10-11 protospacers and one amplicon of approximately 300 bp containing a single protospacer. This allowed us to confirm if the length, position on the genome or protospacer density impacted the effect of eDNA. To our surprise, only the supplementation of a single fragment, Fragment A (Figure 2B) resulted in a significant change in BIM generation (p<0.001, Welch’s Anova, Games-Howell test); a 70(10)% increase. This confirms that epigenetic factors such as DNA modification are not responsible for the effect of eDNA and also indicates that a specific sequence is likely responsible.

### The genomic differences between phage 2972 and phage 858 are clustered

Since the PCR fragment supplementation experiments indicate that epigenetic factors are likely not involved and since only the supplementation of phage 2972 DNA was able to increase BIM generation in a phage 2972 challenge, we surmised that the sequences responsible for this effect are likely located in genomic regions of high variability between phages 2972 and 858. Alignment of the two genomes confirmed that they have 90.2% pairwise identity and indicated not only global homology but also a similar genomic organization. Most of the differences are clustered in a region spanning from 28,505 to 32,484 (positions based on the genome of phage 2972) which covers the phage 2972 ORFs 35 to 38 (phage 858 ORFs 37 to 40) (Figure 3A). A search on NCBI’s conserved domain database showed that nearly all functional domains identified are identical between the four ORFs, indicating a likely shared function despite little pairwise identity at both the nucleotides and amino acid levels. In phage 2972, the region was previously identified as covering the early expressed genes (37) and due to the similarity in annotations between the two phages in it likely that these genes have similar expression patterns in phage 858. Serendipitously, the Fragment A PCR amplicon used earlier happened to be located within this variable region, reinforcing the importance of this region to the effect of eDNA.

**Figure 3:**
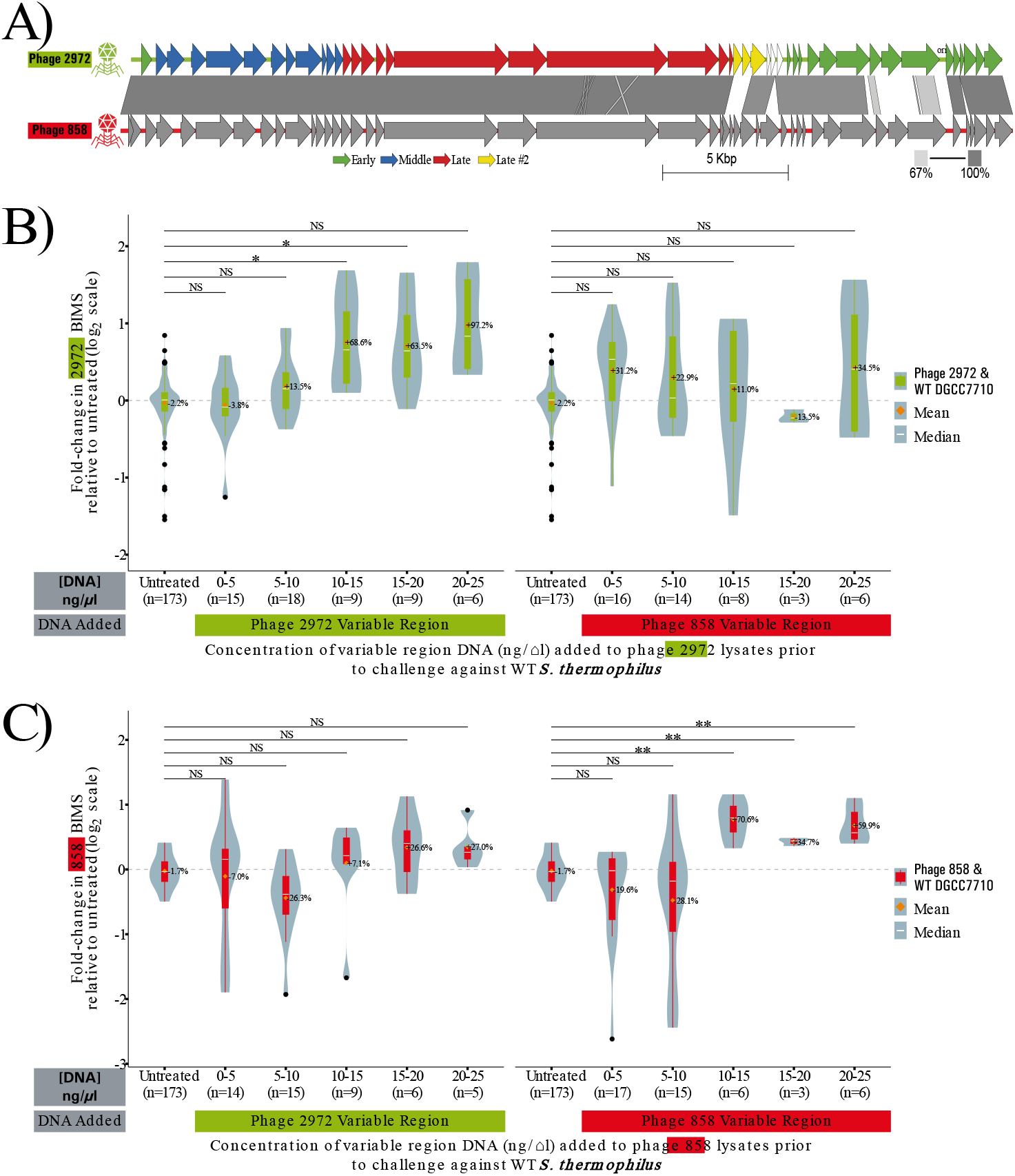
Genomic regions of variability between phages 2972 and 858 increase BIM generation only of the cognate phage. A) Genomic comparison of phages 2972 and 858. CDS are shown as arrows, percent homology by blastn is shown by a gray bar between the genomes, scale from 67% to 100% homology. Transcription modules of phage 2972 according to Duplessis et. al. are shown by colours (green, blue, red and yellow for early, medium, late and late II transcribed genes). The origin of replication for phage 2972 is shown as ori. B-C) BIMs obtained in challenges with phage 2972 (B) and phage 858 (C) lysates supplemented with the variable region of both phages at varying concentrations. Conditions are grouped in bins of concentration of 5 ng/µl. Origin of the variable region is indicated by a coloured bar under the X-axis. BIMs are reported as fold-change relative to the baseline, violin plots represent the probability density curve of the distribution, boxplots represent the first and third quartile of the distribution (box), the minimum and maximum (whiskers), the median (white line) as well as any outliers (black dots). The mean of each distribution is represented by an orange diamond. Significance is determined by pairwise comparison with the untreated condition using Welch’s Anova followed by Games-Howell post-hoc pairwise comparison test.

### A specific region of the phage genome is responsible for the effect of eDNA

To further examine this region, we designed PCR amplicons spanning most of the variable region for both the genomes of phage 2972 and phage 858. We then performed phage challenges using both phage 2972 and phage 858 lysates either untreated or supplemented with various concentrations of variable region PCR amplicon DNA. This was done to test whether the effect of supplemented DNA would vary with the concentration used. In all cases we saw no dose-response effect and only saw an effect when at least 10 ng·µl^-1^ was added. Moreover, there was only a significant effect (p<0.05, Welch’s Anova, Games-Howell test) when the DNA supplemented corresponded to the lysate used in the challenge. In other words, only a phage’s own early region is able to cause an increase in BIM generation (Figure 3B-C).

### eDNA does not serve as a source of genetic material for spacer acquisition

To convincingly show that eDNA provides genetic information for spacer acquisition, we tracked which spacers were acquired by BIMs obtained from lysate supplemented with the PCR fragment A amplicon. Since supplementation of the fragment A amplicon resulted in an increased number of BIMs, if eDNA provides genetic information for spacer acquisition we would expect an over-representation of spacers matching the protospacers present on the amplicon. We sequenced 16 BIMs obtained from the lysate supplemented with fragment A, out of which 12 had acquired a distinct spacer in their CR1 array. When mapping the 12 acquired spacers to the genome of phage 2972, no acquired spacer mapped to the fragment A amplicon. We also sequenced BIMs obtained from other supplemented lysates and for all BIMs where the CRISPR arrays were successfully amplified by PCR we observed acquisition in either the CR1 or CR3 array, consistent with the known dominance of CRISPR-based phage immunity in this system (36).

To verify how likely this scenario was, if eDNA acts as a source of protospacers, we performed a binomial distribution test composed of the following assumptions: 1) eDNA provides genetic material for new spacer acquisition, 2) due to the 70(10)% increase in BIMs when using lysate supplemented with fragment A, 42.7% of the BIMs are due to the effect of eDNA, 3) there is a 3% (7/233 protospacers) change of acquiring a spacer mapping to the amplicon from BIMs not caused by eDNA and 4) there is a 90% chance of acquisition in CR1 (36). This results in a probability of any acquired spacers to map back to Fragment A of 41.1% (90% ∗ (42.7% + 3%)). Using the binomial distribution, we can establish that the probability of observing 0 spacers mapping to Fragment A across 16 BIMs by chance alone is 2.07e-4 which allows us to reject the hypothesis. Since three of our four assumptions are experimentally supported, the most likely faulty assumption is that eDNA provides genetic material for new spacer acquisition.

We arrived at the same conclusion using another experiment based on the search for the acquisition of non-protective spacers. We infected a culture of *S. thermophilus* DGCC7710 in LM17 broth with phage 2972 lysate supplemented with DNA from phage 858. While these two related phages share some protospacers which when acquired would be protective against both phages, they also contain unique protospacers which are only protective against a single phage. We targeted 10 protospacers unique to phage 858 using a multiplex PCR strategy with a shared forward primer binding upstream of the CRISPR array and reverse primers specific to each spacer. As these spacers would not be protective, we extracted the total DNA of the liquid culture (eDNA + bacterial gDNA) to use as a template for the multiplex PCR. In cultures challenged with phage 2972 lysate supplemented with phage 858 DNA, we detected no acquisition of the 10 targeted unique protospacers while they were all detected by gel electrophoresis when tested individually in the culture challenged with phage 858 used as a control. To verify how likely that scenario is, if eDNA acts as a source of protospacers, we performed a binomial distribution test composed of the following assumptions: 1) eDNA provides genetic material for new spacer acquisition, 2) in seven 5 ml cultures at a bacterial concentration established by a growth curve as ∼1e8 CFU/ml and using a CRISPR acquisition rate of ∼1 in 1e6 (20), we can expect 3500 individual BIMs, 3) we can expect 40% of BIMs to be due to eDNA considering the 40(4)% reduction in BIMs caused by DNase treatment, 4) 95.2% of eDNA BIMs are due to the supplemented phage 858 DNA as we supplemented 10 ng/µl of DNA to an average phage eDNA concentration of 0.5(1) ng/µl as established earlier, 5) only 4.3% (10/234) of phage 858’s protospacers are targeted by the multiplex primer mix, and 6) only 25% of the extracted DNA was used for the PCR reaction. This results in a probability of any acquired spacers being one of the 10 targeted spacers of 0.4% (40%*95.2%*4.3%*25%). Using the binomial distribution, we can establish that the probability of not finding any of the targeted spacers across the entire set by chance alone is 6.32e-7 which allows us to reject the hypothesis. Similarly, all but one of our assumptions are experimentally derived and the most likely faulty assumption is the eDNA provides sequence information for CRISPR acquisition. However, this experiment assumes that the DNA extraction process is perfect, and that no DNA is lost. If we account for potential loss of DNA during extraction, even DNA retention rates as low as 22% still yield a probability of randomly not detecting any acquisition below 0.05. While a direct proof of a negative is impossible, these experiments show that eDNA is not likely to provide genetic information for spacer acquisition despite previous work showing that ssDNA can be used as part of the acquisition process (38).

### The effect of eDNA is limited to Oxoid LM17

Throughout the process of performing these experiments, we used LM17 media from the Oxoid (Nepean, Canada) brand. Eventually, Oxoid brand LM17 became unavailable and upon switching to another supplier, we realized that the effect of eDNA on CRISPR adaptation was lost in other brands of LM17. While we were able to generate BIMs using LM17 media obtained from brands Difco (Franklin Lakes, USA),) and Tekniscience (Terrebonne, Canada), there was no longer a significant reduction in BIM generation caused by DNase treatment (Supplementary Figure 5A). Similarly, the effect of eDNA supplementation was also lost in LM17 from Difco and Tekniscience (Supplementary Figure 5B). In an attempt to elucidate this loss of the phenotype, we tested the stability of eDNA and the activity of DNase in LM17 from the various brands as well as in spent cultures in the same media and found that eDNA is stable and DNase is active in all media types.

## Discussion

We show here that eDNA plays a role in the development of immunity against phages (Figure 1) and that it is more specifically phage eDNA matching a specific region corresponding to some of the early expressed genes that is responsible for this effect (Figures 2 and 3). Both removal and supplementation of eDNA in phage lysate influence the amount of BIMs generated, making a clear connection between the naturally present phage eDNA in phage lysate and BIM generation. While the specifics of this interaction remain unknown, we have uncovered some elements required for the effect of eDNA on BIM generation.

The eDNA can be partially degraded, as shown by our restriction enzyme experiments (Supplementary Figure 2), does not need to be modified (Figure 2 b, 3) and only specific regions of the phage genome can affect BIM generation when supplemented (Figure 2b, 3). This indicates a sequence requirement for this effect. The fact that this effect happens in two different phages and that the phage sequences targeted are not cross-reactive (Figure 3) points to a direct sequence recognition of the eDNA to the phage genome rather than a mechanism driven the function of the gene product encoded by the eDNA that is taken up.

Moreover, the loss of the effect of eDNA in a competence deficient background strongly indicates that the eDNA is taken up by the bacteria to mediate this effect (Supplementary Figure 5). However, supplementation of eDNA does not lead to a bias in the acquisition of spacers matching the protospacers present on the supplemented sequences. This shows that while the effect of eDNA is mediated from inside the cell, it does not provide early access to genetic information for spacer acquisition.

Considering this, we propose a model where specific sequences of phage eDNA are taken up by the cell and then interfere with the typical infection cycle – either directly or through some other sequence-dependent anti-phage immune system, allowing more opportunities for the bacteria to acquire spacers through potentially independent mechanisms. The possibility of slowing or blocking phage DNA replication as an adjuvant mechanism of resistance is not unheard of. Previous work has shown that the presence of a phage origin of replication on a plasmid transformed into *S. thermophilus* leads to resistance to phage by preventing the accumulation of phage DNA (39). While the origin of replication of phage 2972 is outside of the variable region PCR amplicon (40) and the Fragment A amplicon, the effect of this region on BIM generation might still be caused by a similar mechanism of stalling phage DNA replication. The early steps of DNA replication involve multiple protein-ssDNA interactions (41, 42), all of which could be impacted by competitive binding to “dummy sequences” acquired from eDNA. Further experiments would need to be done to confirm the internal mechanism of the acquired eDNA and its potential interactions with the CRISPR-Cas system and without media from the Oxoid supplier they are current impossible. Overall, our data link the presence of phage eDNA and CRISPR adaptation. As a natural byproduct of phage-derived lysis, the presence of phage eDNA is already associated with the threat of lysis for bacteria and its involvement in the development of CRISPR immunity shows that bacteria have multiple ways of being safely exposed to phage DNA. Moreover, this discovery brings another tool in the phage-bacteria interaction toolbox, providing the community with another way to influence BIM generation, by modulating the levels of phage eDNA present in lysates. This can open up new avenues of research into the mechanism of CRISPR acquisition and potentially lead to the discovery of new factors influencing its activity during phage infection.

## Methods

### Strains used and culturing

*S. thermophilus* DGCC7710 was grown in LM17 media composed of M17 powder (unless otherwise specified Oxoid, Nepean, Canada) supplemented with 5 g·L^-1^ without agitation at 37°C for overnight cultures and at 42°C for cultures used for same-day experiments (day cultures). Lysates of phages 2972 and 858 (40) were amplified and quantified as described (20) in LM17-CaCl (LM17 media supplemented with 10 mM of CaCl_2_).

### Purification of phage particles

To purify phage particles, 0.5 M of NaCl and 10% w·v^-1^ of PEG8000 were added to 1 L of phage lysate and incubated overnight at 4°C with gentle agitation. The phages were precipitated by centrifugation at 20,000 g for 15 min at 4°C and the pellet was resuspended in 5 ml of phage buffer. The solution was centrifuged at 210,000 g for 3 h on a CsCl gradient column formed of three 8 ml solutions of CsCl at concentrations of 0.525, 0.775 and 1.025 g·ml-1 respectively (lowest on top). Fractions of approximately 500 μl were collected starting at the indicative blue diffraction band. All fractions were tested for phage titre and the fractions containing phage were also tested for the presence of DNA by treating a sample of each fraction with MseI and HindII (New England Biolabs, Whitby, Canada) restriction enzymes for 15 min at 37°C according to the manufacturer’s recommendations. The samples were then run on a 1% agarose gel to confirm the absence of contaminant genomic DNA (Not shown).

### BIM assays

Phage lysate containing between 1e8 and 3e8 PFU*ml^-1^ were diluted to 80% original concentration with a combination of phage buffer (50 mM Tris-HCl, pH 7.5, 100 mM NaCl, 8 mM MgSO4), purified DNA, DNase buffer, DNase or other supplemented enzymes or protein depending on the condition. For the MOI range assays, phage lysates were diluted with phage buffer only to the desired MOI. For the assay, 100 µl of diluted phage and 300 µl of bacterial culture at OD_600_ between 0.4 and 0.5 as measured on a Genesys 30 Spectrophotometer (Thermo Scientific, Waltham, USA) were added to 3 ml of molten LM17-CaCl 0.75% w·v^-1^ agar and poured over LM17 1% w ·v^-1^ agar plates. The plates were incubated for 24-48h at 42°C in sealed plastic bags before the colonies were counted. CFU counts were analyzed using the R Statistical Software (v4.2.2) (43). For most BIM assays, in order to account for the inherent variability in BIMs obtained from different bacterial cultures on different days, BIMs were reported as the log2 of the count relative to the baseline. The baseline was established for each day’s bacterial culture and phage lysate pair used by averaging the CFUs obtained from the untreated condition.

When required, single colonies were picked and used as a template for amplification by PCR using Q5 DNA polymerase (New England Biolabs, Whitby, Canada) with primers CR1_F and R as well as CR3_F and R (Supplementary Table 2) according to the manufacturer’s recommendation. After verification of the size of the amplicons to confirm acquisition by gel electrophoresis, the PCR amplicons were purified used the Monarch PCR and DNA Cleanup Kit (New England Biolabs, Whitby, Canada) and sent for Sanger sequencing at the McMaster Genomics facility. The newly acquired spacers were identified by direct sequence alignment.

### DNA extraction from phage particles

Lysates were treated with 20 μg·ml_-1_ of RNase A and 2U·ml_-1_ of DNase I (New England Biolabs, Whitby, Canada) along with 10% DNase buffer 10x at 37°C for 30 min followed by heat inactivation at 75°C for 10 min. The phages were then treated with 100 μg·ml_-1_ of Proteinase K (New England Biolabs, Whitby, Canada) with SDS to a final concentration of 2% (v·v^-1^) at 37°C for 1 h. DNA was extracted using phenol:chloroform (1:1), precipitated using an isopropyl alcohol wash and ethanol and then rehydrated in Elution Buffer (New England Biolabs, Whitby, Canada). DNA concentration was measured using a QuBit 4 Fluorometer and the QuBit dsDNA High Sensitivity Quantitation kit (Invitrogen, Waltham, USA) and DNA purity was assessed using a DS-11 series spectrophotometer (DeNovix, Wilmington, USA).

### DNA extraction from bacteria

Bacterial cultures were grown overnight and pelleted by centrifugation for 1-5 min at >12 000 g and resuspended in 0.8% of the original volume of 10 mM Tris-Cl pH 8.0. DNA was extracted using the Monarch Genomic DNA Purification Kit (New England Biolabs, Whitby, Canada) according to the manufacturer’s recommendations for DNA purification from gram-positive bacteria and Archaea. DNA concentration was measured using a QuBit 4 Fluorometer and the QuBit dsDNA High Sensitivity Quantitation kit (Invitrogen, Waltham, USA) and DNA purity was assessed using a DS-11 series Spectrophotometer (DeNovix, Wilmington, USA).

### Digestion of eDNA in lysates

To fully digest eDNA from phage lysates, 256 μl of phage lysate was added to 32 μl of DNase Buffer or rCutSmart Buffer (for restriction enzymes) along with 10 μl of either DNase I, MseI, HindIII-HF or NotI-HF (New England Biolabs, Whitby, Canada) and 22 μl of phage buffer. The mixture was incubated at 37°C for 1 h and then kept at 4°C for no more than 2 h before being used for a BIM generation assay. Controls treated with buffer only were also performed in a similar manner by replacing the volume of enzyme with phage buffer.

### Quantification of eDNA by qPCR

Quantification of eDNA was performed by qPCR using the Azure Cielo real-time PCR machine (Azure Biosystems, Dublin, USA) and primers specific to phage 858, phage 2972 and to S. thermophilus DGCC7710. The reaction was performed using the PowerUp SYBR Green Master Mix (Applied Biosystems, Waltham, USA) according to the manufacturer’s recommendations using the manufacturer’s standard cycling mode for primers with T_m_ > 60°C. Standard curves were established with all three primer sets by performing qPCR on serial dilutions of a solution containing a known quantity (measured using Fluorometer and the QuBit dsDNA High Sensitivity Quantitation kit (Invitrogen, Waltham, USA) of phage 858, phage 2972 or S. thermophilus DGCC7710 DNA. Data analysis was performed using the R Statistical Software (v4.2.2) (43). The Ct values measured (n=3) were correlated with the known concentration of DNA and a linear regression model was established using the stats R package (v4.0.3) (R Core Team, 2022). The model was used to convert measured Ct values into DNA concentrations and standard error was carried along using the errors R package (v0.4.0) (44).

### Liquid BIM assays and multiplex PCR

Bacterial cultures were grown to an OD_600_ of 0.4 and 5 ml of culture was added to 1.33 ml of phage lysate diluted to arrive at a final MOI of 0.3. CaCl_2_ was added to a final concentration of 10 mM. Control cultures were supplemented with phage buffer instead of phage lysate. The cultures were incubated overnight at 42°C and the cells were then pelleted and the gDNA was extracted. From 3 ml of the supernatant, eDNA was extracted using phenol:chloroform:isoamyl alcohol (25:24:1, v·v^-1^), precipitated using an isopropyl alcohol wash and ethanol and then rehydrated in Elution Buffer (New England Biolabs, Whitby, Canada).

The extracted DNA was then used as the template in a PCR reaction with a set of primers designed to target 10 spacers matching unique protospacers to 858. The primers were designed to avoid potential off target binding to other protospacers, from either phage 2972 or phage 858. The PCR reaction was set up using Q5 polymerase (New England Biolabs, Whitby, Canada) following the manufacturer’s recommendation, using an annealing temperature of 51.4°C for CR1 and 52.7°C for CR3, chosen as the average recommended annealing temperature of all the primer pairs in the multiplex primer set. Results were visualized using agarose gel electrophoresis where the presence of a band indicates acquisition of a spacer matching one of the 10 targeted protospacers. The ability for the multiplex PCR to readily detect acquisition was confirmed by PCR on liquid culture infected with phage 858.

### Construction of the ΔcomEC strain

The pCR-ComEC plasmid containing 1) the pNZ123 backbone, 2) an artificial mini-CRISPR array composed of the CR1 leader sequence and of a single spacer targeting the comEC gene flanked by two repeats and 3) a recombination template composed of 500 bp upstream and downstream of the comEC gene was assembled using Gibson Assembly (New England Biolabs, Whitby, Canada) according to the manufacturer’s recommendation. The resulting plasmid was transformed into commercial NEB5α E. coli cells, according to the manufacturer’s recommendations (New England Biolabs, Whitby, Canada). The plasmid was then purified using the NEB Miniprep kit and then electroporated into electrocompetent *S. thermophilus* DGCC7710 made through a glycine shock protocol as described in (20).

Colonies of *S. thermophilus* growing on LM17 supplemented with 30 μg·μl-1 of chloramphenicol (Cm30) were picked and tested by PCR to show deletion of the comEC gene. The confirmed mutant was then serially passaged on LM17 media without antibiotics to cure the pCR-ComEC plasmid. Loss of the plasmid was assessed by a loss of resistance to chloramphenicol in liquid media by inoculating a single colony into two LM17 tubes, only one of which contained chloramphenicol.

## Supplementary

**Supplementary Figure 1:**
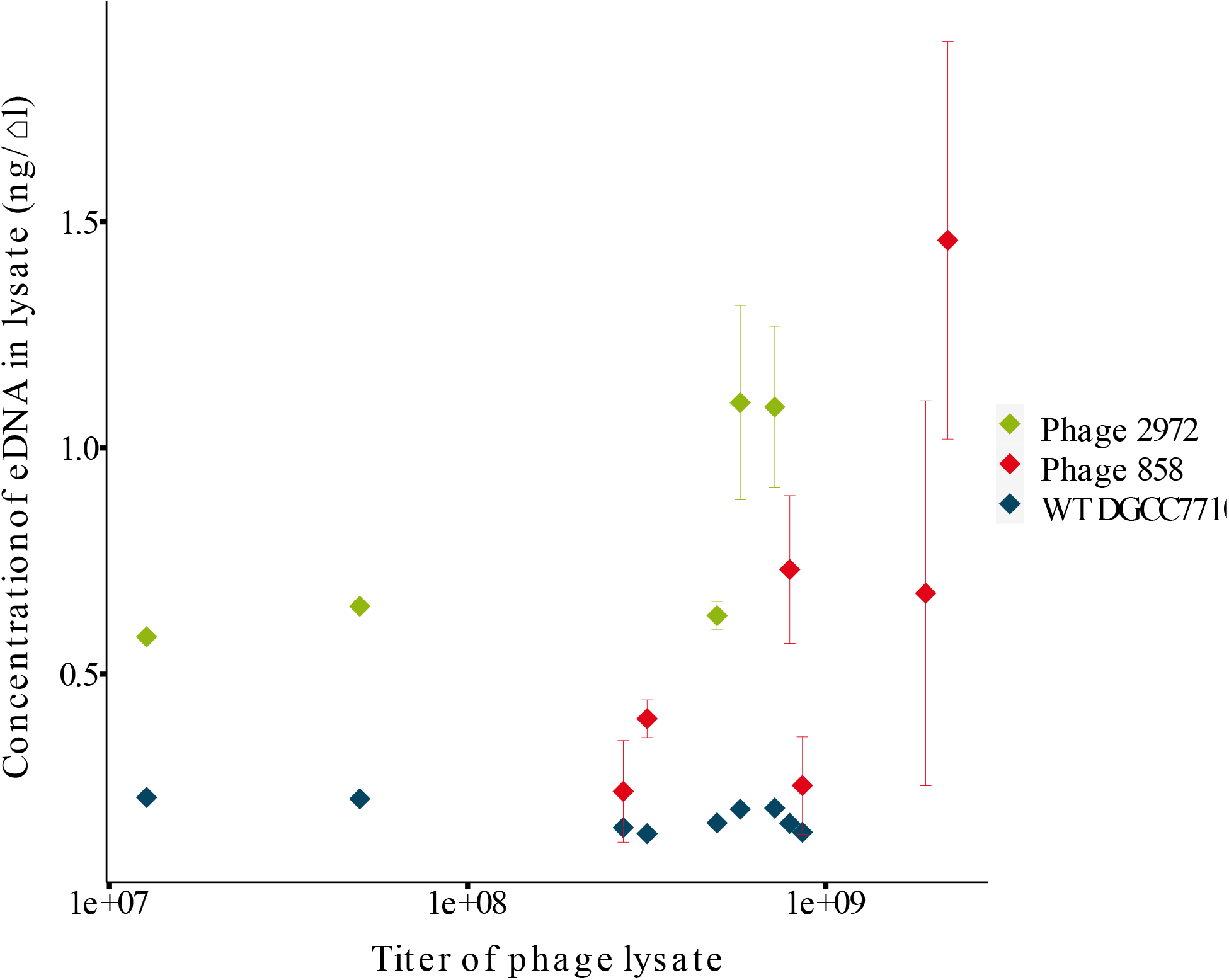
Concentration of eDNA as a function of phage titre. Phage 2972 values are shown in green, phage 858 values in red and DGCC7710 in blue. Linear regression analysis showed no correlation between phage titre and concentration of eDNA both when considering each phage alone and when considering all lysates together. Error bars represent the standard error on each value carried over from the linear regression model established from the qPCR standard curve. Standard error was carried using the first-order Taylor series method. Two phage 858 samples were never tested with the WT DGCC7710 primers.

**Supplementary Figure 2:**
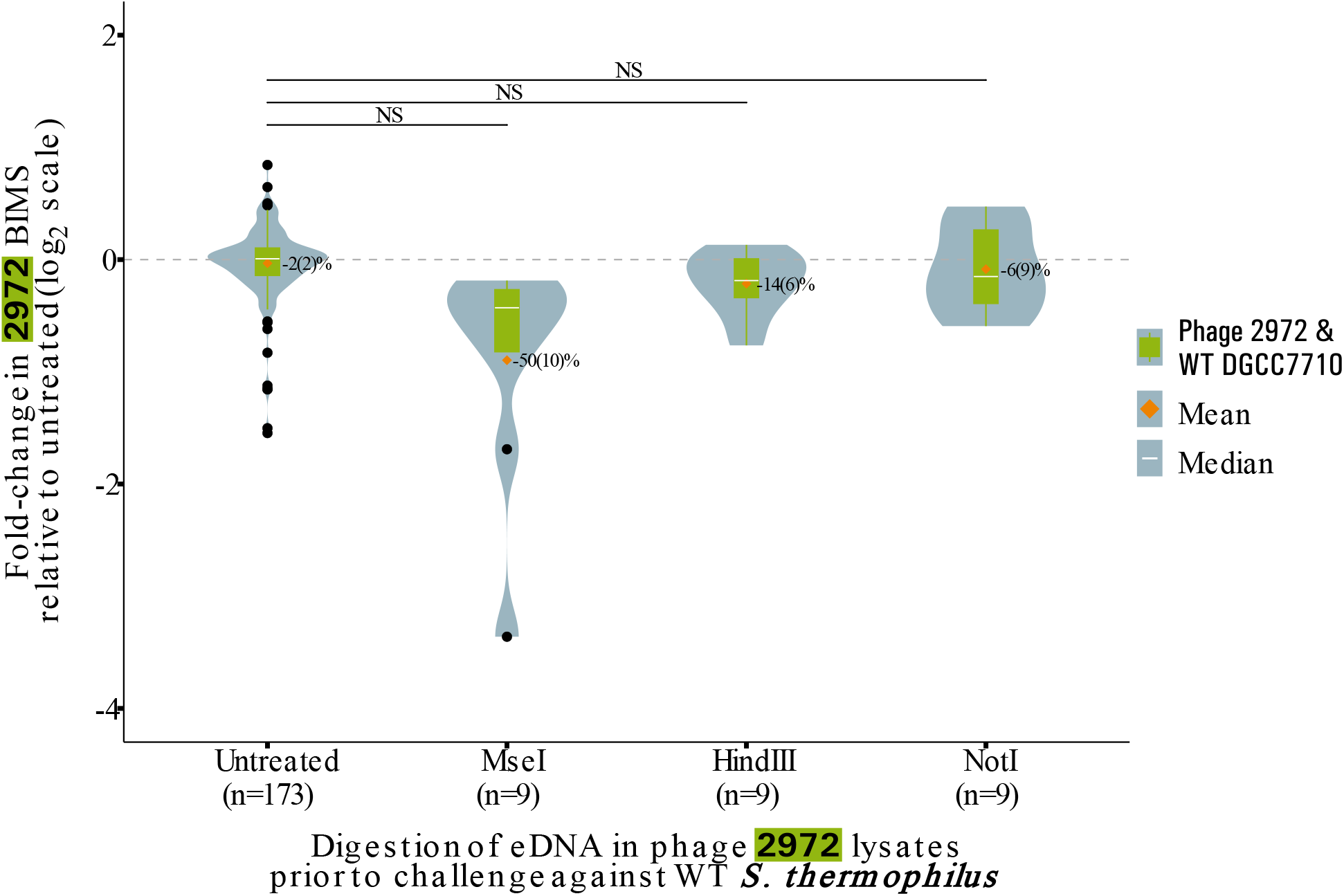
BIM generation with lysate treated with restriction enzymes. BIMs obtained in challenges against phage 2972. BIMs are reported as fold-change relative to the baseline, violin plots represent the probability density curve of the distribution, boxplots represent the first and third quartile of the distribution (box), the minimum and maximum (whiskers), the median (white line) as well as any outliers (black dots). The mean of each distribution is represented by an orange diamond. Significance is determined by pairwise comparison with the untreated condition using Welch’s Anova followed by Games-Howell post-hoc pairwise comparison test. The restrictions enzymes are predicted to cut the genome of phage 2972 269 times for MseI, 16 times for HindIII and 0 times for NotI.

**Supplementary Figure 3:**
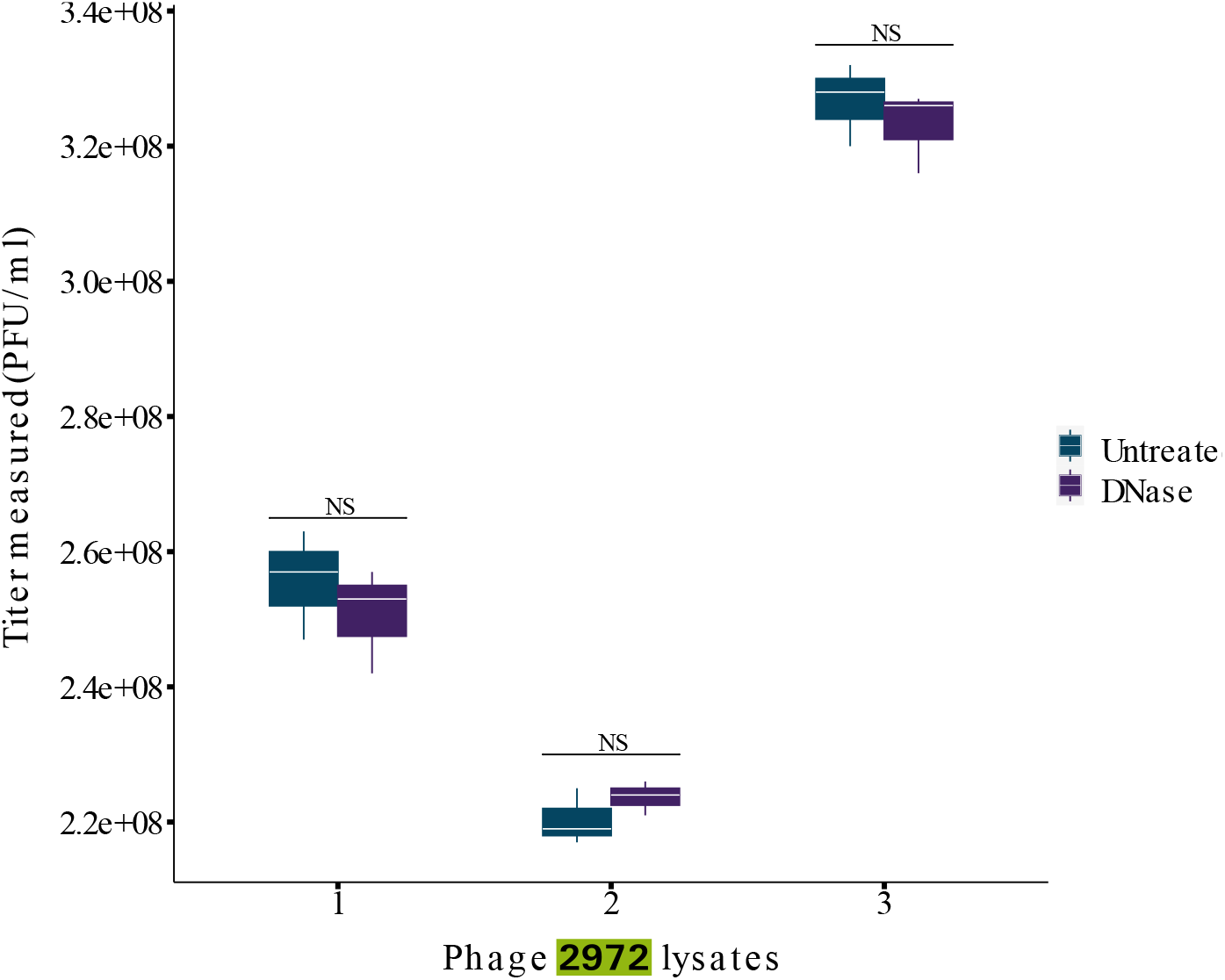
Effect of DNase treatment on phage titer. Titer of various phage 2972 lysates before and after DNase treatment. Each lysate was quantified using a full plate PFU assay with 3 replicates for each measurement. The DNase treatment did not significantly impact the titer of phage 2972 lysates according to the Tukey HSD test (p>0.05).

**Supplementary Figure 4:**
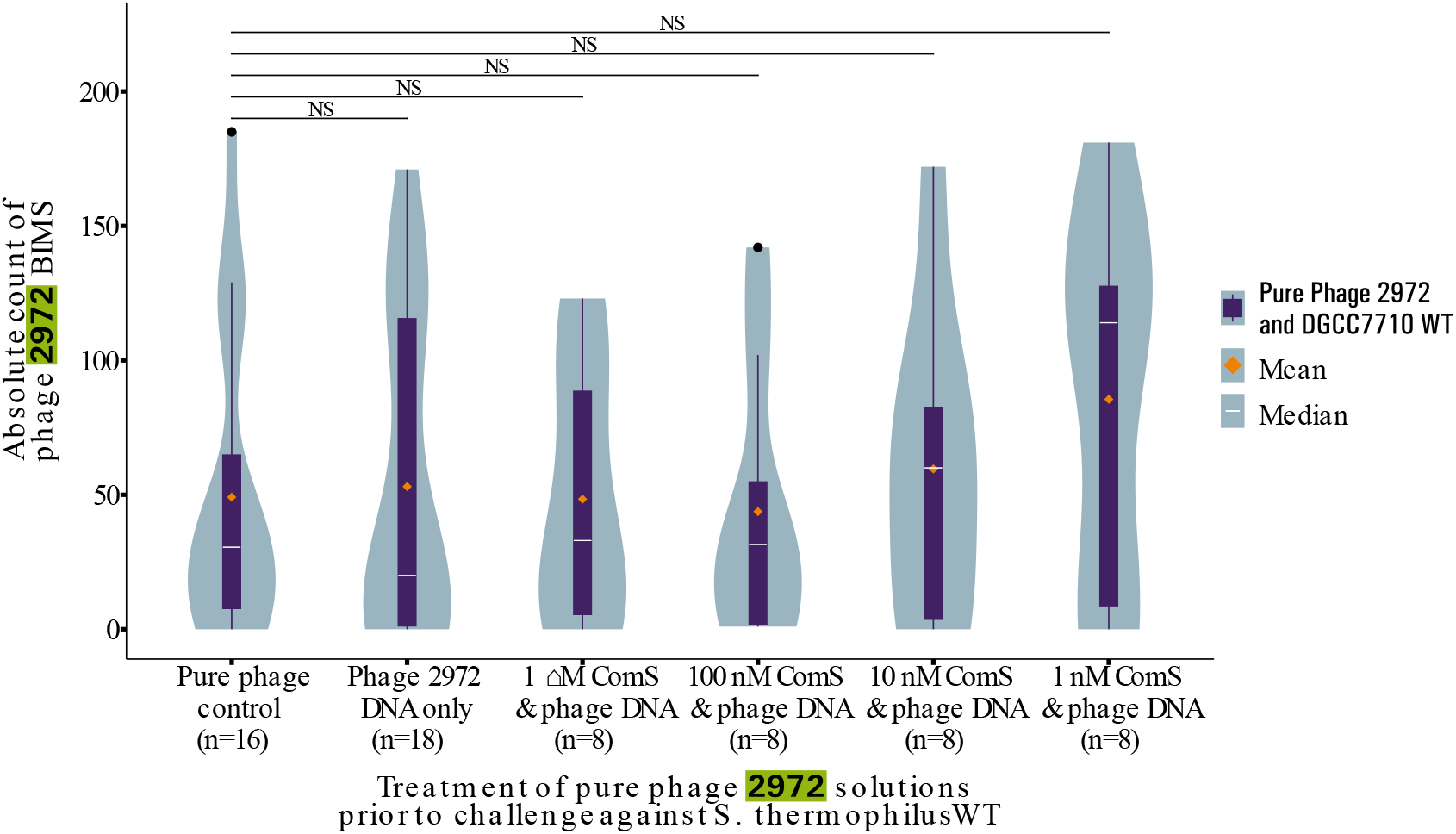
BIM generation with purified phage supplemented with phage DNA and ComS peptide. BIMs obtained in challenges against phage 2972. BIMs are reported as absolute CFU count, violin plots represent the probability density curve of the distribution, boxplots represent the first and third quartile of the distribution (box), the minimum and maximum (whiskers), the median (white line) as well as any outliers (black dots). The mean of each distribution is represented by an orange diamond. Significance is determined by pairwise comparison with the untreated condition using Welch’s Anova followed by Games-Howell post-hoc pairwise comparison test. When supplemented, phage DNA was supplemented to a final concentration of 10 ng·μl^-1^.

**Supplementary Figure 5:**
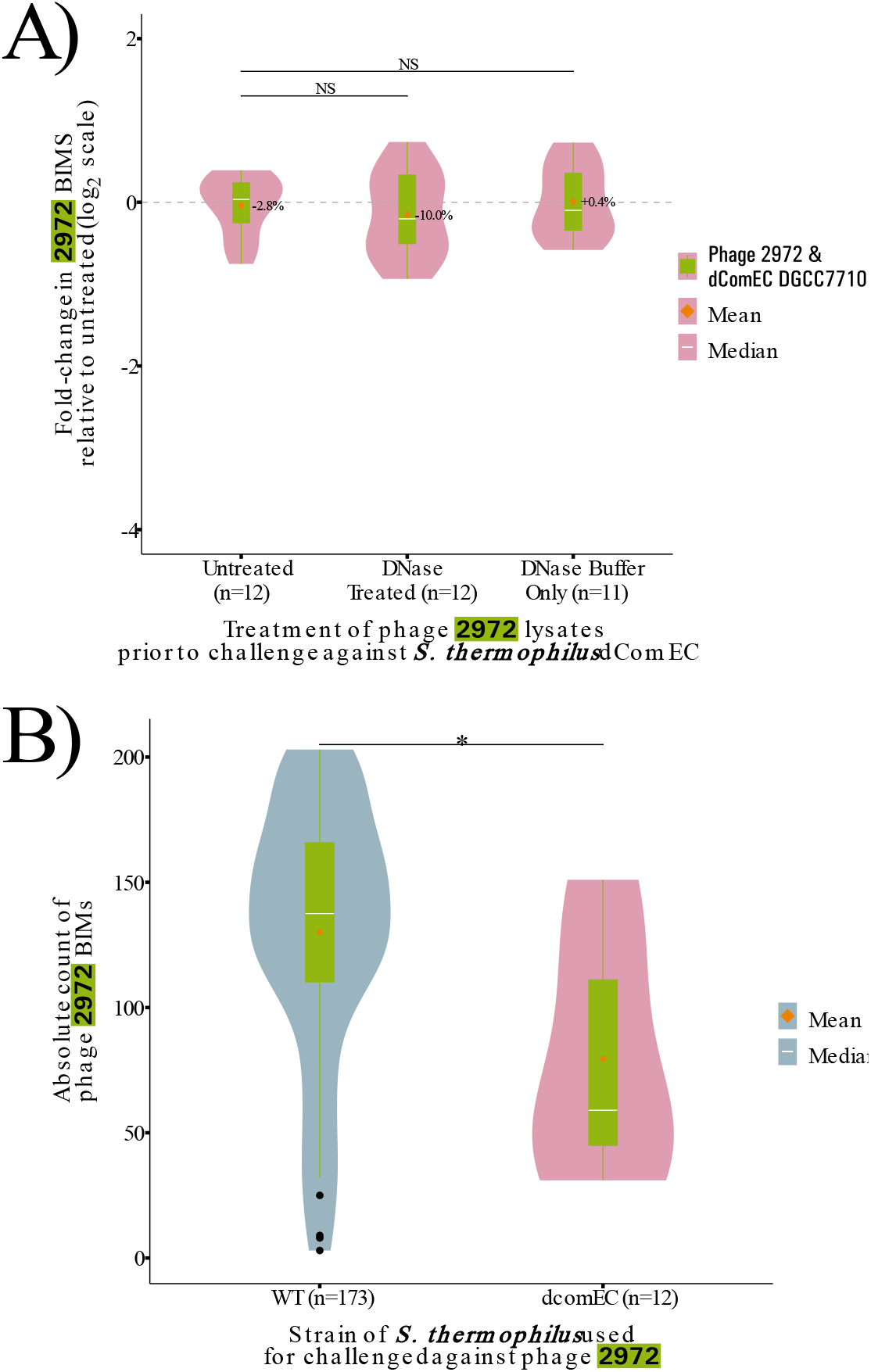
Difference in effect of eDNA in *S. thermophilus* WT vs ΔcomEC. A) BIMs obtained in challenges of phage 2972 against *S. thermophilus* ΔcomEC. BIMs are reported as fold-change relative to the baseline. B) BIMs obtained in challenges of phage 2972 against *S. thermophilus* WT and ΔcomEC. BIMs are reported as absolute CFU count. In all cases, violin plots represent the probability density curve of the distribution, boxplots represent the first and third quartile of the distribution (box), the minimum and maximum (whiskers), the median (white line) as well as any outliers (black dots). The mean of each distribution is represented by an orange diamond. Significance is determined by pairwise comparison with the untreated condition using Welch’s Anova followed by Games-Howell post-hoc pairwise comparison test.

**Supplementary Figure 6:**
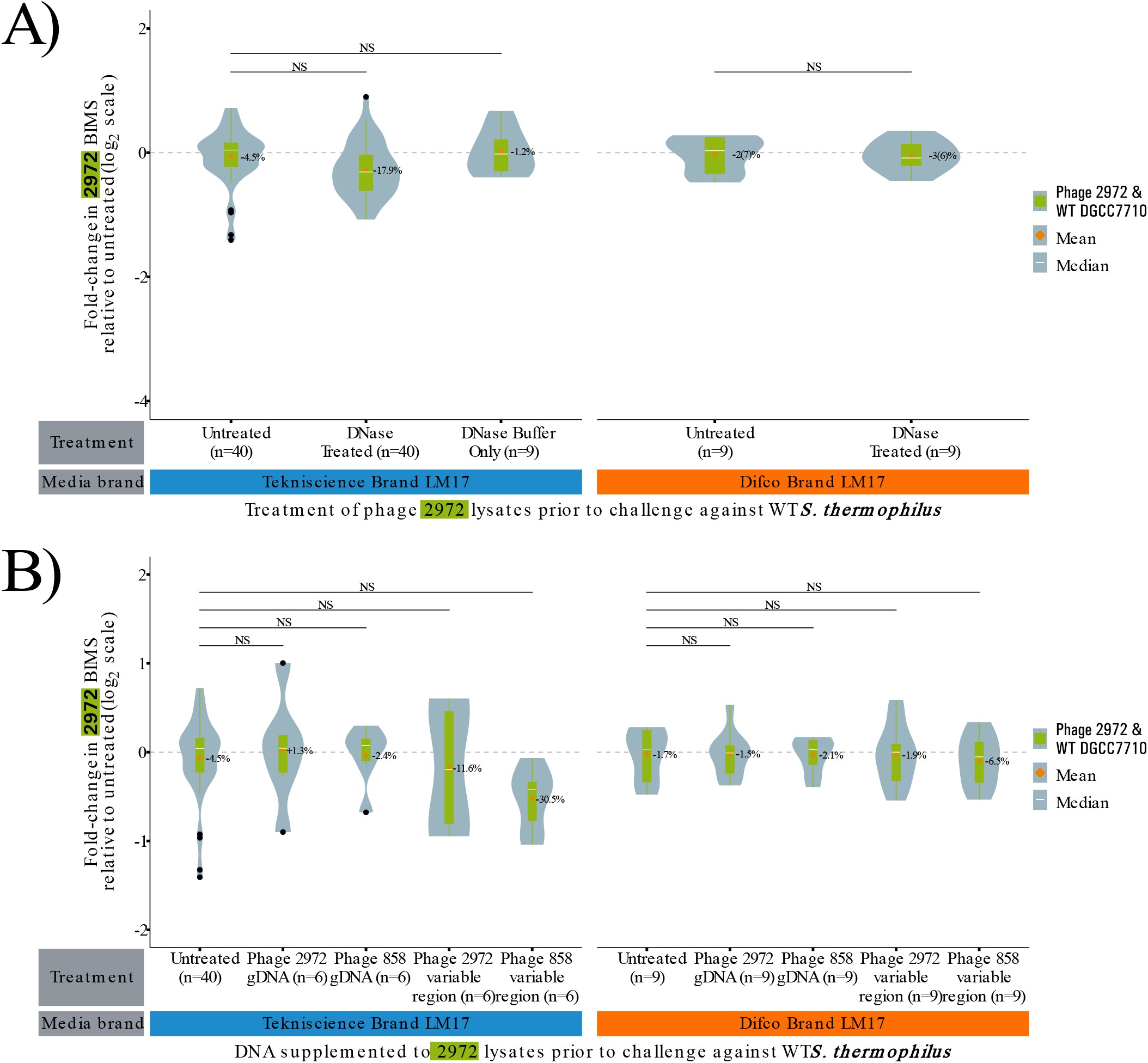
BIM assay in LM17 from different brands. A) BIMs obtained in challenges with phage 2972 lysates treated with DNase in Tekniscience and Difco brand LM17. B) BIMs obtained in challenges with phage 2972 lysates supplemented to a final concentration of 10 ng·μl^-1^ with varying DNA sources in Tekniscience and Difco brand LM17. In all cases, violin plots represent the probability density curve of the distribution, boxplots represent the first and third quartile of the distribution (box), the minimum and maximum (whiskers), the median (white line) as well as any outliers (black dots). The mean of each distribution is represented by an orange diamond. Significance is determined by pairwise comparison with the untreated condition using Welch’s Anova followed by Games-Howell post-hoc pairwise comparison test. The media brand used is indicated by a coloured bar under the X-axis.

**Supplementary Table 1:** Strains and plasmids used.

| Name | Source | Reference |
| --- | --- | --- |
| *S. thermophilus* DGCC7710 | Félix d'Hérelle Phage Reference Center | HER 1458 |
| Brussowvirus bv2972 |  | HER 458 |
| Brussowvirus bv858 |  | HER 459 |
| pNZ123 |  | (45) |

**Supplementary Table 2:**
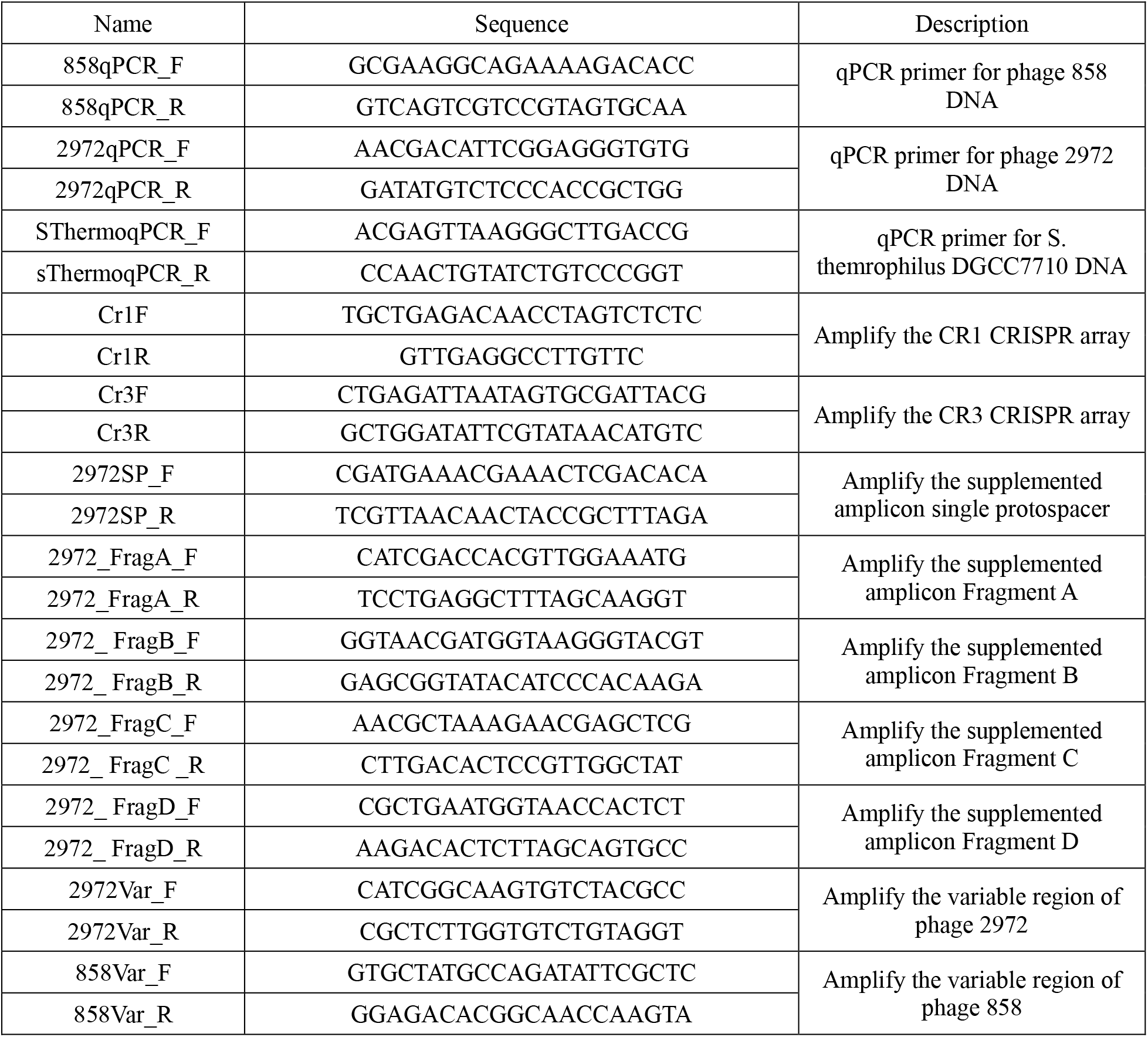
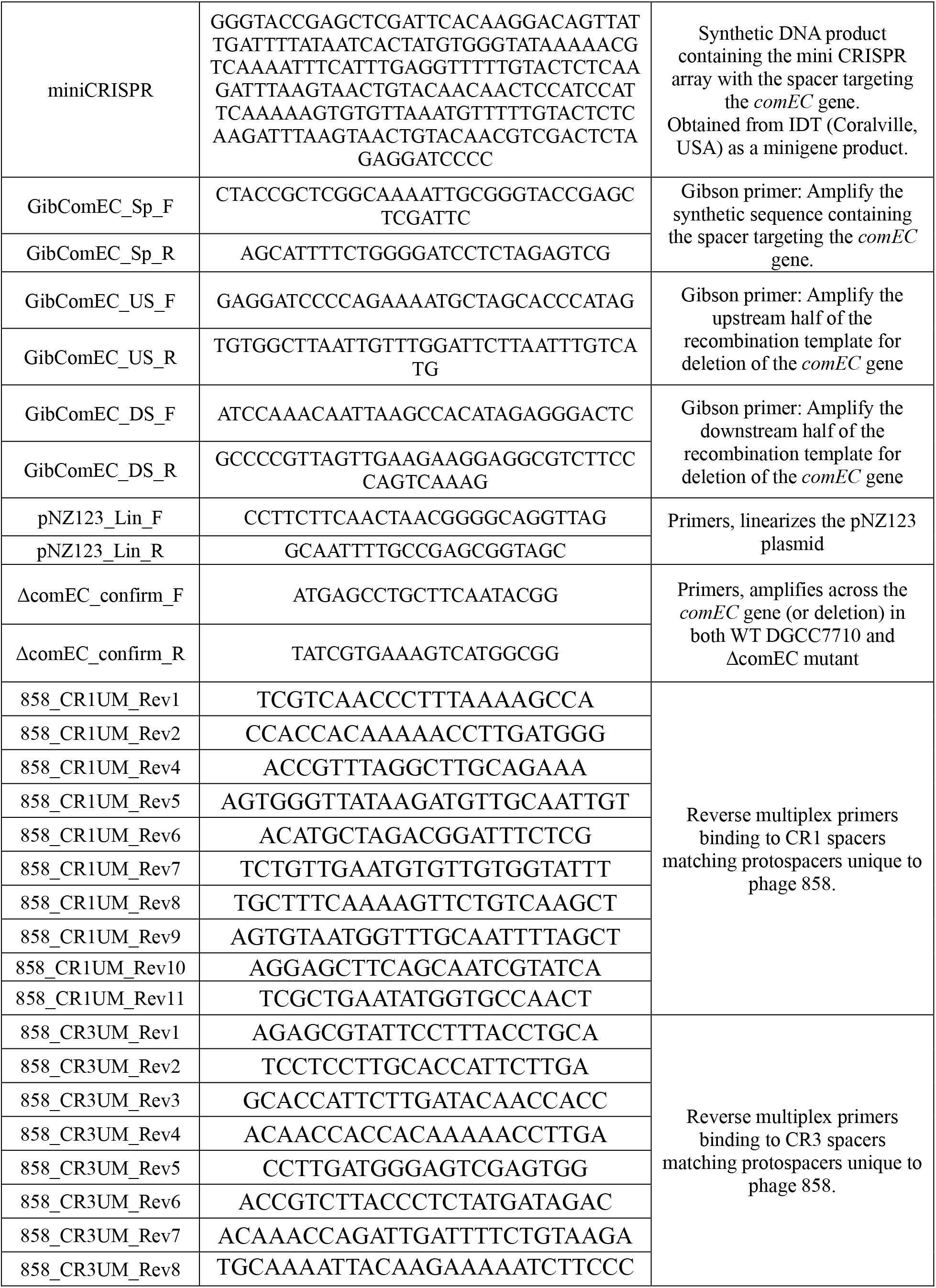
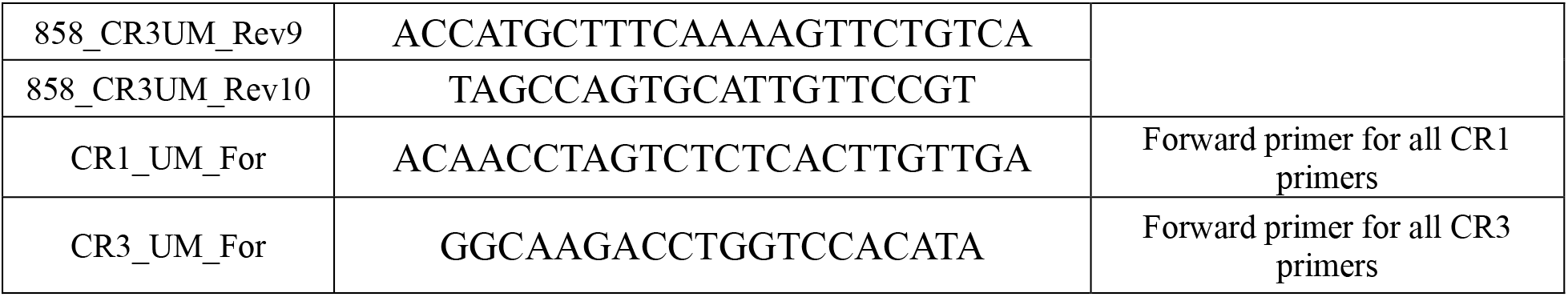
DNA products used.

